# Geometric constraints dominate the antigenic evolution of influenza H3N2 hemagglutinin

**DOI:** 10.1101/014183

**Authors:** Austin G. Meyer, Claus O. Wilke

## Abstract

We have carried out a comprehensive analysis of the determinants of human influenza A H3 hemagglutinin evolution, considering three distinct predictors of evolutionary variation at individual sites: solvent accessibility (as a proxy for protein fold stability and/or conservation), experimental epitope sites (as a proxy for host immune bias), and proximity to the receptor-binding region (as a proxy for protein function). We found that these three predictors individually explain approximately 15% of the variation in site-wise *dN*/*dS*. The solvent accessibility and proximity predictors were largely independent of each other, while the epitope sites were not. In combination, solvent accessibility and proximity explained 32% of the variation in *dN*/*dS*. Incorporating experimental epitope sites into the model added only an additional 2 percentage points. We also found that the historical H3 epitope sites, which date back to the 1980s and 1990s, showed only weak overlap with the latest experimental epitope data. Finally, sites with *dN*/*dS* > 1, i.e., the sites most likely driving seasonal immune escape, are not correctly predicted by either historical or experimental epitope sites, but only by proximity to the receptor-binding region. In summary, proximity to the receptor-binding region, and not host immune bias, seems to be the primary determinant of H3 evolution.

**Author summary:** The influenza virus is one of the most rapidly evolving human viruses. Every year, it accumulates mutations that allow it to evade the host immune response of previously infected individuals. Which sites in the virus’ genome allow this immune escape and the manner of escape is not entirely understood, but conventional wisdom states that specific “immune epitope sites” in the protein hemagglutinin are preferentially attacked by host antibodies and that these sites mutate to directly avoid host recognition; as a result, these sites are commonly targeted by vaccine development efforts. Here, we combine influenza hemagglutinin sequence data, protein structural information, experimental immune epitope data, and historical epitopes to demonstrate that neither the historical epitope groups nor epitopes based on experimental data are crucial for predicting the rate of influenza evolution. Instead, we find that a simple geometrical model works best: sites that are closest to the location where the virus binds the human receptor are the primary driver of hemagglutinin evolution. There are two possible explanations for this result. First, the existing historical and experimental epitope sites may not be the real antigenic sites in hemagglutinin. Second, alternatively, hemagglutinin antigenicity may not the primary driver of influenza evolution.

## Introduction

The influenza virus causes one of the most common infections in the human population. The success of influenza is largely driven by the virus’s ability to rapidly adapt to its host and escape host immunity. The antibody response to the influenza virus is determined by the surface proteins hemagglutinin (HA) and neuraminidase (NA). Among these two proteins, hemagglutinin, the viral protein responsible for receptor binding and uptake, is a major driver of host immune escape by the virus. Previous work on hemagglutinin evolution has shown that the protein evolves episodically [1–3]. During most seasons, hemagglutinin experiences mostly neutral drift around the center of an antigenic sequence cluster; in those seasons, it can be neutralized by similar though not identical antibodies, and all of the strains lie near each other in antigenic space [4–7]. After several seasons, the virus escapes its local sequence cluster to establish a new center in antigenic space [7–9].

There is a long tradition of research aimed at identifying important regions of the hemagglutinin protein, and by proxy, the sites that determine sequence-cluster transitions [4, 6, 10–21]. Initial attempts to identify and categorize important sites of H3 hemagglutinin were primarily sequence-based and focused on substitutions that took place between 1968, the emergence of the Hong Kong H3N2 strain, and 1977 [10, 11]. Those early studies used the contemporaneously solved protein crystal structure, a very small set of mouse monoclonal antibodies, and largely depended on chemical intuition to identify antigenically relevant amino-acid changes in the mature protein. Many of the sites identified in those studies reappeared nearly two decades later, in 1999, as putative epitope sites with no additional citations linking them to actual immune data [4]. Those sites and their groupings are still considered the canonical immune epitope set today [3, 16, 22]. While the limitations of experimental techniques and of available sequence data in the early 1980’s made it necessary to form hypotheses based on chemical intuition, these limitations are starting to be overcome through recent advances in experimental immunological techniques and wide-spread sequencing of viral genomes. Therefore, it is time to revisit the question of whether or not the host immune system directly pressures influenza to evolve to escape antibody binding, or perhaps, there is some other indirect manner of immune escape. For example, at least one recent model has suggested that the hemagglutinin protein may evolve to modulate receptor-binding avidity rather than to modulate antibody-binding [23]. Moreoever, since the original epitope set was identified via sequence analysis, we do not even know whether *bona fide* immune-epitope sites actually exist, i.e., sites which represent a measurable bias in the host immune response. Most importantly, even if immune-epitope sites do exist and can be experimentally identified, it is possible that they do not experience more positive selection than other important sites in the protein.

Some recent studies have begun to address these questions indirectly, via evolutionary analysis. For example, over the last two decades, virtually every major study on positive selection in hemagglutinin has found some but never all of the historical epitope sites to be under positive selection [3, 16, 18, 19, 23]. Furthermore, each of these studies has found a set of sites that are under positive selection but do not belong to any historical epitope. Finally, because every study identifies slightly different sites, there seems to be no broad agreement on which sites are under positive selection [12,16,18,19]. The sites found by disparate techniques are similar but they are never identical.

To dissect the determinants of hemagglutinin evolution, we here linked several predictors, including relative solvent accessibility, the inverse distance from the receptor-binding region, and experimental immune epitope data, to site-wise evolutionary rates calculated from all of the human H3N2 sequence data for the last 22 seasons (1991–2014). We found that, individually, all these predictors explained approximately 15% of evolutionary rate variation. In addition, we analyzed all of the available H3 experimental epitope data, and we found that current experimental data does not at all reflect the historical epitope sites or their groups. After controlling for biophysical constraints with relative solvent accessibility and function with distance to the receptor-binding region, the remaining predictive power of either experimental or historical categories was relatively low. Finally, by explicitly accounting for RSA, proximity, and host immune data, we found that we could predict nearly 35% of the evolutionary rate variation in hemagglutinin, nearly twice as much variation as could be explained by earlier models.

## Results

### Relationship between evolutionary rate and inverse distance to the receptor-binding site

Our overarching goal in this study was to identify specific biophysical or biochemical properties of the mature protein that determine whether a given site will evolve rapidly or not. As a measure of evolutionary variation and selective pressure, we used the metric *dN*/*dS*. *dN*/*dS* can measure both the amount of purifying selection acting on a site (when *dN*/*dS* ≪ 1 at that site) and the amount of positive diversifying selection acting on a site (when *dN*/*dS* ≳ 1). For simplicity, we will refer to *dN*/*dS* as an *evolutionary rate*, even though technically it is a *relative* evolutionary rate or evolutionary-rate ratio. We built an alignment of 3854 full-length H3 sequences spanning 22 seasons, from 1991/92 to 2013/14. We subsequently calculated *dN*/*dS* at each site, using a one-rate fixed-effects likelihood (FEL) model as implemented in the software HyPhy [24].

Several recent works have shown that site-specific evolutionary variation is partially predicted by a site’s solvent exposure and/or number of residue-residue contacts in the 3D structure [19, 20, 25–30] (see Ref. [31] for a recent review). This relationship between protein structure and evolutionary conservation likely reflects the requirement for proper and stable protein folding: Mutations at buried sites or sites with many contacts are more likely to disrupt the protein’s conformation [30] or thermodynamic stability [32]. In addition, there may be functional constraints on site evolution. For example, regions in proteins involved in protein–protein interactions or enzymatic reactions are frequently more conserved than other regions [27, 33, 34]. However, these structural and functional constraints generally predict the amount of purifying selection expected at sites, and therefore they cannot identify sites under positive diversifying selection. Moreover, the short divergence time of viruses causes the systematic biophysical pressures that predict much of eukaryotic protein evolution to be much less dominant in viral evolution [28]. Thus, we set out to find a constraint on hemagglutinin evolution that was related to the protein’s role in viral binding and fusion.

A few earlier studies had shown that sites near the sialic acid-binding region of hemagglutinin tend to evolve more rapidly than the average for the protein [4, 20, 21]. Furthermore, when mapping evolutionary rates onto the hemagglutinin structure, we noticed that the density of rapidly evolving sites seemed to increase somewhat towards the receptor-binding region (Fig. 1A). Therefore, as the primary function of hemagglutinin is to bind to sialic acid and induce influenza uptake, we reasoned that distance from the receptor-binding region of HA might serve as a predictor of functionally driven HA evolution. We calculated distances from the sialic acid-binding region (defined as the distance from site 224 in HA), and correlated these distances with the evolutionary rates at all sites. We found that distance from the receptor-binding region was a strong predictor of evolutionary rate variation in hemagglutinin (Pearson correlation *r* = 0.41, *P* < 10^−15^).

**Figure 1.**
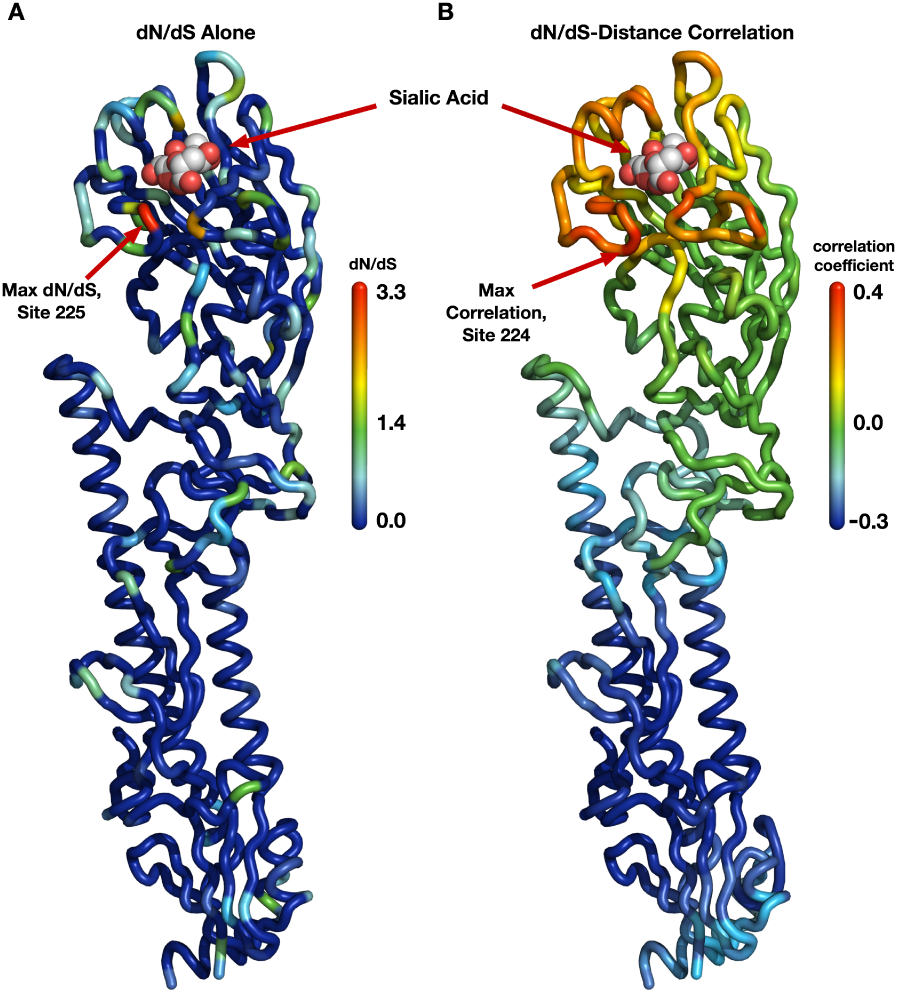
Evolutionary-rate variation along the hemagglutinin structure. (A) Each site in the protein structure is colored according to its evolutionary rate *dN*/*dS*. Hot colors represent high *dN*/*dS* (positive selection) while cool colors represent low *dN*/*dS* (purifying selection). (B) Each site in the protein structure is colored according to the *dN*/*dS*–distance correlation obtained when distances are calculated relative to that site. Hot colors represent positive correlations while cool colors represent negative correlations. Thus, distances from sites that are redder are better positive predictors of the evolutionary rates in the protein than are distances from bluer sites; distances from blue sites are actually anti-correlated with evolutionary rate. Distances from sites that are colored green have essentially no predictive ability.

Next, we wanted to verify that this correlation was representative of hemagglutinin evolution and not just an artifact of the specific site chosen as the reference point in the distance calculations. It would be possible, for example, that distances to several spatially separated reference sites all resulted in similarly strong correlations. We addressed this question systematically by making, in turn, each individual site in HA the reference site, calculating distances from that site to all other sites, and correlating these distances with evolutionary rate. We then mapped these correlations onto the structure of hemagglutinin, coloring each site according to the strength of the correlation we obtained when we used that site as reference in the distance calculation (Fig. 1B). We obtained a clean, gradient-like pattern: The correlations were highest when we calculated distances relative to sites near the receptor-binding site (with the maximum correlation obtained for distances relative to site 224), and they continuously declined and then turned negative the further we moved the reference site away from the apical region of hemagglutinin (Fig. 1B). This result was in stark contrast to the pattern we had previously observed when mapping evolutionary rate directly (Fig. 1A). In that earlier case, while there was a perceptible preference of faster evolving sites to fall near the receptor-binding site, the overall distribution of evolutionary rates along the structure looked mostly random to the naked eye. We thus found a geometrical, distance-based constraint on hemagglutinin evolution: Sites evolve faster the closer they lie toward the receptor-binding region.

We also evaluated how proximity to the receptor-binding region performed as a predictor of *dN*/*dS* in comparison to the previously proposed structural predictors relative solvent accessibility (RSA) and weighted contact number (WCN). We found that among these three quantities, proximity to the sialic acid-binding region was the strongest predictor, explaining 16% of the variation in *dN*/*dS* (Pearson *r* = 0.41, *P* < 10^−15^, see also Figs. 2 and S1). RSA and WCN explained 14% and 6% of the variation in *dN*/*dS*, respectively (*r* = 0.37, *P* < 10^−15^ and *r* = 0.25, *P* = 7 × 10^−9^). Proximity to the sialic acid-binding region and RSA were virtually uncorrelated (*r* = 0.08, *P* = 0.09) while RSA and WCN correlated strongly (*r* = −0.64, *P* < 10^−15^). These results suggested that proximity to the sialic acid-binding region and RSA should be used jointly in a predictive model.

**Figure 2.**
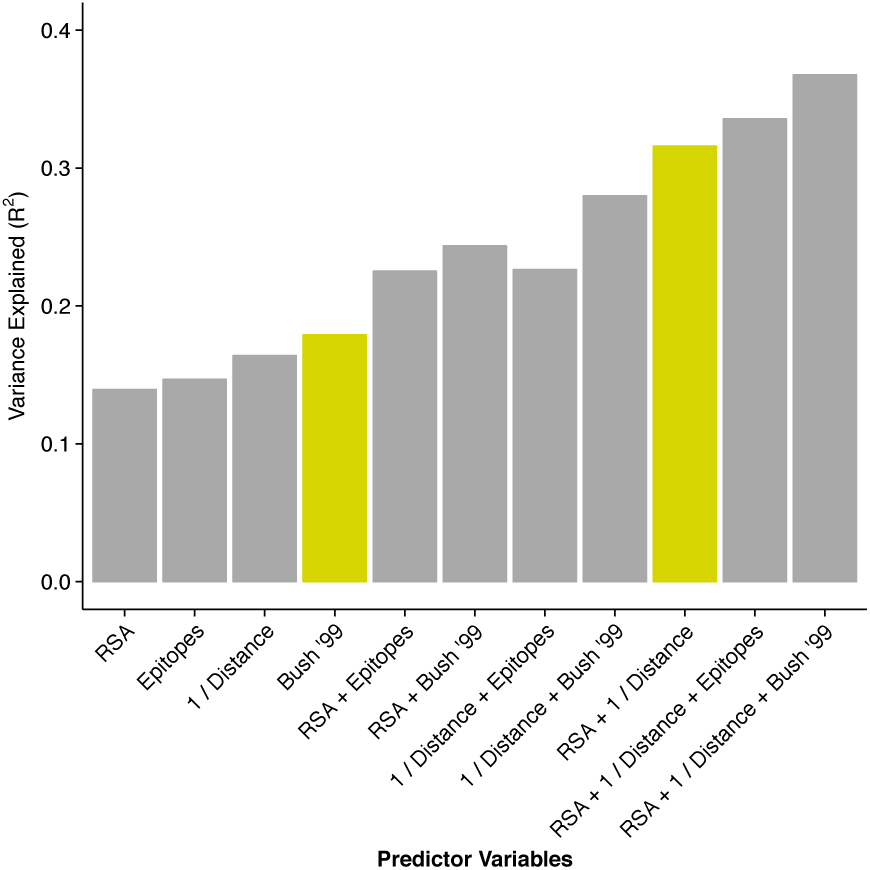
Proportion of variance in *dN*/*dS* explained by different linear models. The height of each bar represents the coefficient of determination (*R*^2^) for a linear model consisting of the stated predictor variables. The historical epitope sites from Bush 1999 [4] (yellow bar on the left) are the single best predictor of evolutionary rate variation. However, a model using two predictors that each have a clear biophysical meaning (solvent exposure, proximity to receptor-binding region) explains almost twice the variation in *dN*/*dS* (yellow bar on the right).

**Figure S1:**
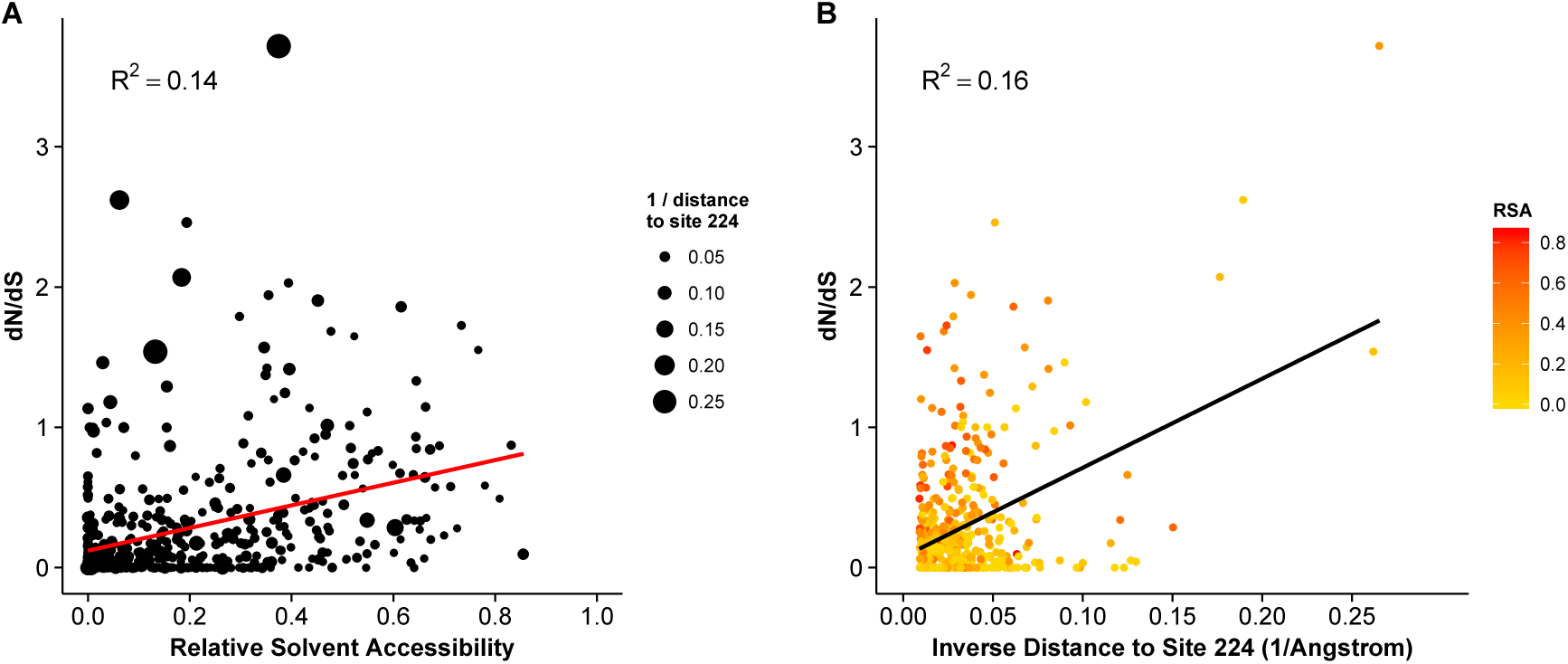
Dependence of *dN*/*dS* on solvent exposure and proximity to the receptor-binding region. (A) *dN*/*dS* vs. RSA. The size of the dots represents 1/Distance. (B) *dN*/*dS* vs. 1/Distance. The coloring of the dots represents RSA. The distance to the sialic acid-binding region is the single strongest quantitative predictor of evolutionary rate ratio in hemagglutinin.

Because hemagglutinin has, in addition to its function as a receptor-binding protein, a host of other intermediate functional states during the viral fusion process, we also tested the ability of structural metrics from the post-fusion state to predict hemagglutinin evolutionary rate [35]. We found no significant metric, either RSA or proximity, derived from the post-fusion state. (Complete data and analysis scripts are available in the accompanying github repository, see Methods for details.)

### Incorporating experimental immunological data

Another potential functional constraint on hemagglutinin evolution is a bias in the human immune system. This bias, generally referred to as antigenicity, describes the extent to which the human immune system does a better job attacking one region of a protein compared to another. Conventional wisdom states that functionally important sites in the protein that are targeted by antibodies will evolve more rapidly to facilitate immune escape. And indeed, our results from the previous subsection have shown that proximity to the receptor-binding region is a good predictor of evolutionary variation. However, if substitutions to avoid direct antibody binding are the primary cause of positive selection, then we would expect antigenic sites on hemaggalutinin to serve as a substantially better predictors of adaptation than proximity to the receptor-binding site alone.

For influenza hemagglutinin H3, there exists a list of canonical, historical epitope sites that are commonly considered to represent this bias [4]. However, these sites were not primarily defined based on actual immunological data, and they have not been re-validated since the late 1990s even though more experimental data is now available. (See Discussion for details on the history of the historical epitope sites.) Before we could generate a combined evolutionary model, we therefore considered it essential to validate the antigenic groups with available immunological data. As it turns out, the majority of antigenic data available did not agree with the historical epitope sites (Supporting Text S1). Therefore, we used both the historical epitope sites and a set of experimentally re-defined epitopes for further modeling.

A detailed explanation of our re-grouping based on experimental data is available in the Supplementary Text S1. It is important to note that these groups are not intended to represent a new canonical set of hemagglutinin epitopes. Indeed, the data from which they were derived is limited and relatively poorly annotated. However, considering the magnitude of the difference between the historical epitopes and the available experimental data we considered it imperative to include experimentally derived epitopes in our analysis.

Thus, we considered both the historical epitope groups (Bush 1999) and the experimentally derived epitopes 1–4, defined in the Supplementary Text. Because a site’s epitope status is a categorical variable, we calculated variance explained as the coefficient of determination (*R*^2^) in a linear model with *dN*/*dS* as the response variable and epitope status as the predictor variable. We found that experimental epitopes explained 15% of the variation in *dN*/*dS*, comparable to RSA and proximity. In comparison, the historical epitopes alone explained nearly 18% of the variation in *dN*/*dS*, outperforming all other individual predictor variables considered here (Fig. 2 and Table 1). However, as discussed in the Supplementary Text S1, the available experimental data suggest that not all of the historical sites may be actual immune epitope sites. Therefore, we suspected that some of the predictive power of historical sites was due to these sites simply being solvent-exposed sites near the receptor-binding region. We similarly wondered to what extent the predictive power of the experimental epitope sites was attributable to the same cause, since, in fact, both historical and experimental epitope sites showed comparable enrichment in sites near the sialic acid-binding region and in solvent-exposed sites (Fig. S2). Therefore, we analyzed how the variance explained increased as we combined epitope sites (experimental or historical) with either RSA or proximity or both.

**Table 1.**
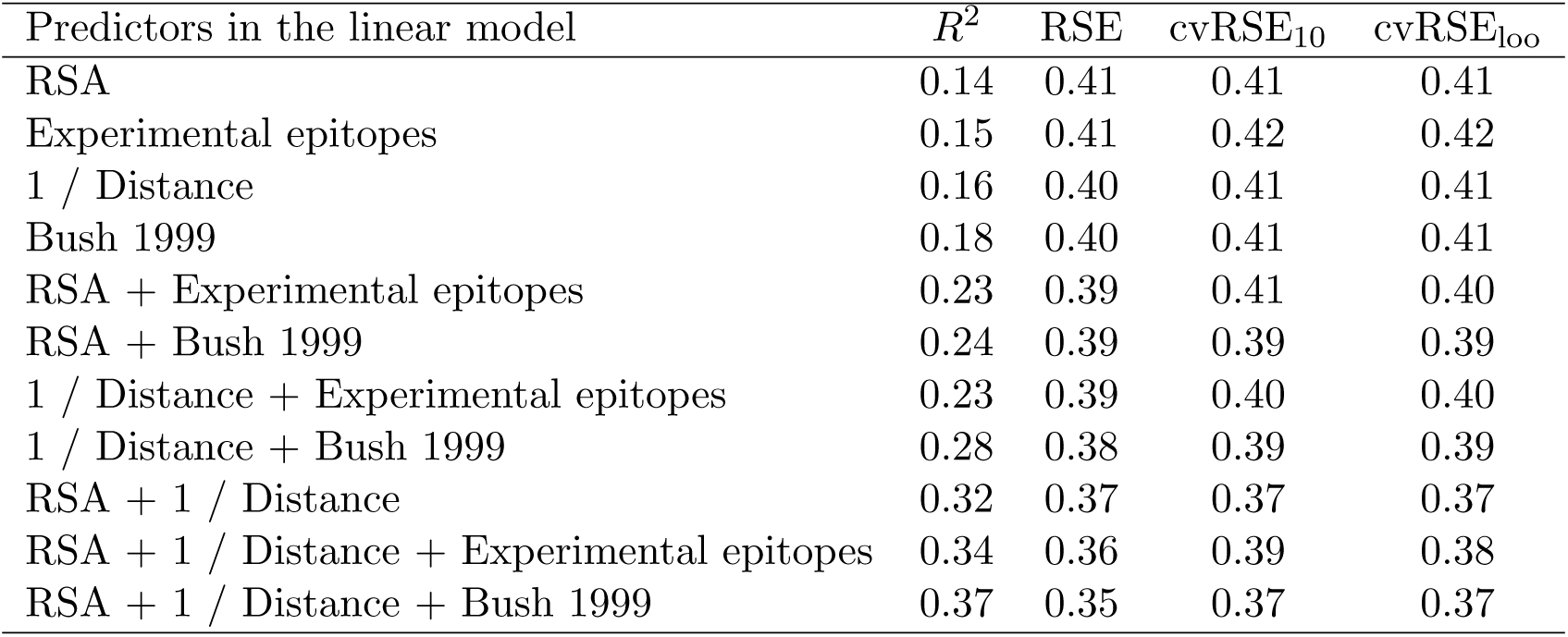
Predictive performance of each linear model considered. *R*^2^ is the proportion of variation in *dN*/*dS* explained by the specified model. RSE is the residual standard error of the linear model. cvRSE_10_ is the cross validated residual standard error calculated by 10-fold cross validation. cvRSE_loo_ is the cross validated residual standard error calculated by leave-one-out cross validation.

| Predictors in the linear model | $R^2$ | RSE | $\text{cvRSE}_{10}$ | $\text{cvRSE}_{\text{loo}}$ |
| --- | --- | --- | --- | --- |
| RSA | 0.14 | 0.41 | 0.41 | 0.41 |
| Experimental epitopes | 0.15 | 0.41 | 0.42 | 0.42 |
| 1 / Distance | 0.16 | 0.40 | 0.41 | 0.41 |
| Bush 1999 | 0.18 | 0.40 | 0.41 | 0.41 |
| RSA + Experimental epitopes | 0.23 | 0.39 | 0.41 | 0.40 |
| RSA + Bush 1999 | 0.24 | 0.39 | 0.39 | 0.39 |
| 1 / Distance + Experimental epitopes | 0.23 | 0.39 | 0.40 | 0.40 |
| 1 / Distance + Bush 1999 | 0.28 | 0.38 | 0.39 | 0.39 |
| RSA + 1 / Distance | 0.32 | 0.37 | 0.37 | 0.37 |
| RSA + 1 / Distance + Experimental epitopes | 0.34 | 0.36 | 0.39 | 0.38 |
| RSA + 1 / Distance + Bush 1999 | 0.37 | 0.35 | 0.37 | 0.37 |

**Figure S2:**
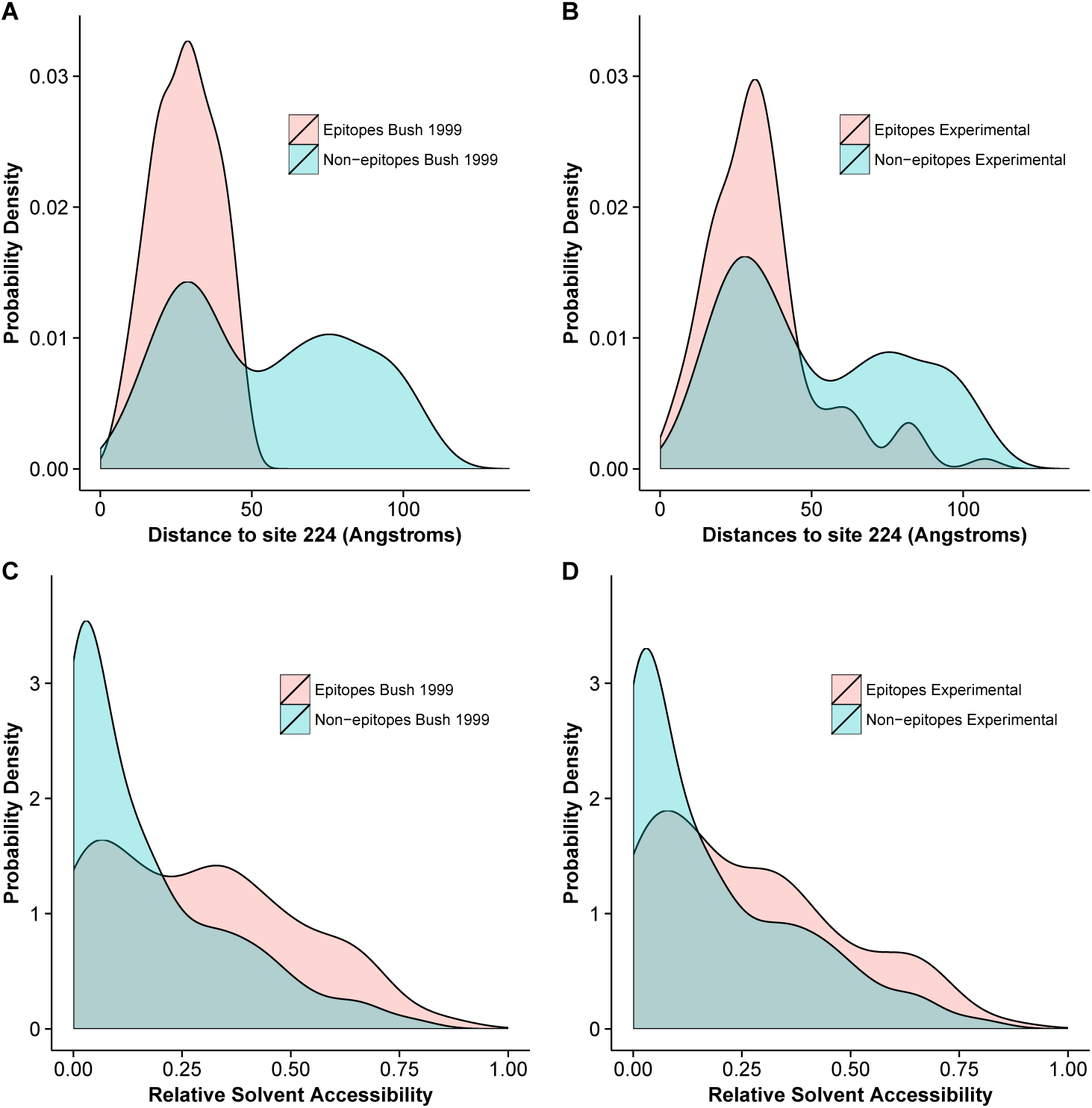
Distance to receptor-binding site and solvent exposure for epitope and non-epitope sites. (A) Distribution of distances to residue 224, for historical epitope and nonepitope sites. (B) Distribution of distances to residue 224, for experimental non-linear epitope and non-epitope sites. (C) Distribution of relative solvent accessibilities, for historical epitope and non-epitope sites. (D) Distribution of relative solvent accessibilities, for experimental nonlinear epitope and non-epitope sites. Under both historical and experimental epitope definitions, epitope sites are closer to the sialic acid-binding region and have higher RSA than non-epitope sites.

We found that epitope status, under either definition (experimental/historical), led to increased predictive power of the model when combined with either RSA or proximity (Fig. 2). However, a model consisting of just the two predictors RSA and proximity, not including any information about epitope status of any sites, performed even better than any of the other one-or two-predictor models, explaining 32% of the variation in *dN*/*dS* (Fig. 2). Adding epitope status to this best-performing two-predictor model resulted in only minor improvement, from 32% to 34% variance explained in the case of experimental epitopes and from 32% to 37% variance explained in the case of historical epitope sites (Fig. 2 and Table 1).

### Predicting sites under selection and comparisons to other work

The geometrical constraints RSA and proximity explained more variance in *dN*/*dS* than did epitope sites, but were they also better at predicting sites of interest? Because *dN*/*dS* can measure purifying as well as positive diversifying selection, the percent variance in *dN*/*dS* that a model explains may not necessarily accurately reflect how useful that model is in predicting specific sites, e.g. sites under positive selection. For example, one could imagine a scenario in which a model does exceptionally well on sites under purifying selection (*dN*/*dS* ≪ 1) but fails entirely on sites under positive selection (*dN*/*dS* > 1). Such a model might explain a large proportion of variance but be considered less useful than a model that overall predicts less variation in *dN*/*dS* but accurately pinpoints site under positive selection. Therefore, we wondered whether epitope sites might do a poor job predicting background purifying selection but might still be useful in predicting sites with *dN*/*dS* > 1. We found, to the contrary, that neither the historical nor the experimental epitope sites could reliably predict sites with *dN*/*dS* > 1, alone or in combination with RSA (Fig. 3A–D). Proximity to the receptor-binding site, on the other hand, correctly predicted four sites with *dN*/*dS* > 1, even in the absence of any other predictors. Notably, all models we considered here were robust to cross-validation. The cross-validated residual standard error was virtually unchanged from its non-cross-validated value in all cases (Table 1). Because proximity clearly identified four points with high *dN*/*dS*, we also verified that the proximity–*dN*/*dS* correlation was not caused just by these four points. We removed from our data set the four points that had both predicted and observed *dN*/*dS* > 1, and found that a significant proximity–*dN*/*dS* correlation remained nonetheless (*r* = 0.17, *p* = 0.00001).

**Figure 3.**
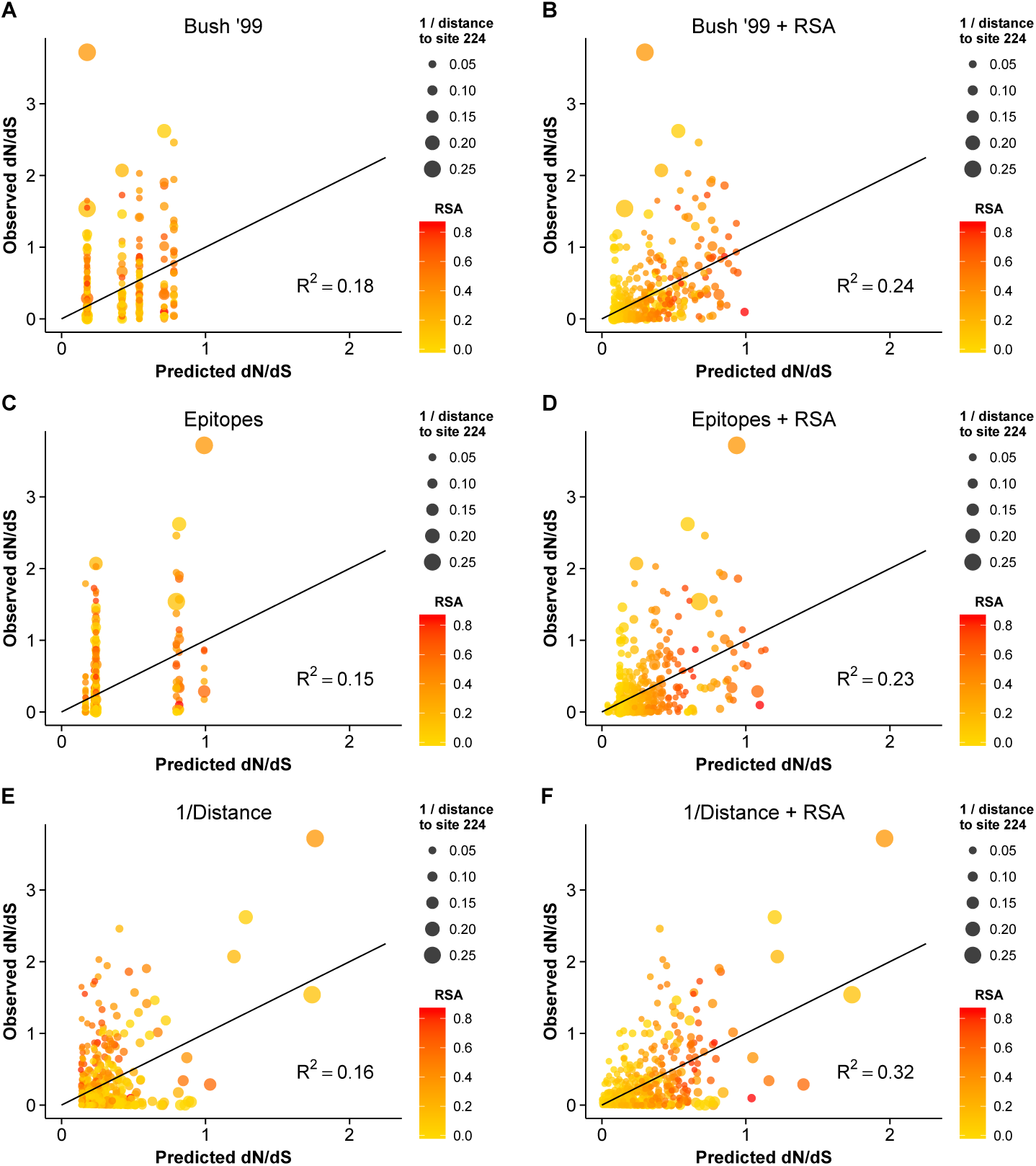
Observed *dN*/*dS* vs. predicted *dN*/*dS* for different predictive linear models. (A) Only epitope status according to the historical definition is used as predictor variable. (B) Historical epitope sites and RSA are used as predictor variables. (C) Only epitope status according to the experimental non-linear epitope data is used as predictor variable. (D) Experimental epitope sites and RSA are used as predictor variables. (E) Only proximity to the sialic acid-binding region (measured as 1/Distance to Residue 224) is used as predictor variable. (F) Proximity and RSA are used as predictor variables. Individual sites with *dN*/*dS* > 1 are predicted correctly only if the linear model includes the 1/Distance predictor. However, in all cases, adding the RSA predictor significantly improves the model predictions.

Finally, we compared the predictions from the geometrical model of hemagglutinin evolution to results from a recent study of antigenic cluster transitions; that study found seven sites near the receptor-binding region which were critical for cluster transitions according to hemagglutinin inhibition (HI) assays with ferret antisera [21]. The sites identified in Ref. [21] were 145, 155, 156, 158, 159, 189, and 193. For comparison, our geometric model (with predictors RSA and 1/Distance) predicted none of these sites to be under positive selection. Sites predicted to have *dN*/*dS* > 1 were instead 96, 137, 138, 143, 222, 223, 225, and 226. Moreover, out of the seven sites from Ref. [21], only one (site 145) had an observed *dN*/*dS* significantly above 1. By contrast, four of the eight sites predicted under the geometric model to have *dN*/*dS* > 1 did indeed have *dN*/*dS* significantly above 1. Thus, the sites that determine the major antigenic changes in the virus did not at all overlap with the sites expected and observed to be under the greatest evolutionary pressure. When investigating the location of these sites in detail, we found that all of the sites we predicted to have *dN*/*dS* > 1 were located just basal to the receptor-binding site, whereas nearly all of the sites from [21] (with the exception of 145, the site with *dN*/*dS* > 1) were located on the apical side of the receptor-binding site (Fig. 4).

**Figure 4.**
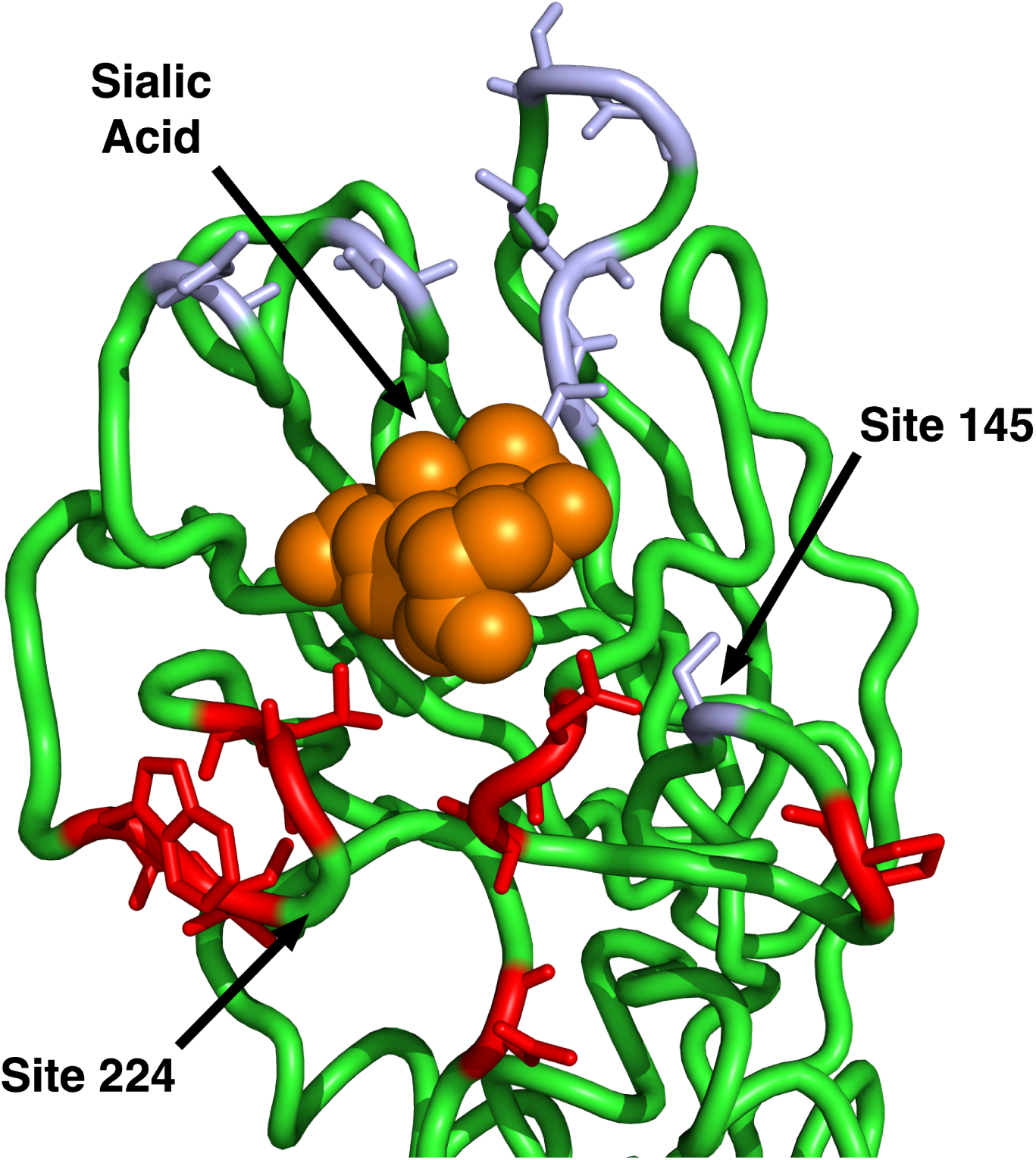
**Sites identified by Koel et al. 2013 and those predicted to have** *dN*/*dS* > 1. The sites shown in purple are those identified by Koel et al. 2013 [21] to be critical for antigenic cluster transitions. Only one of these sites has a *dN*/*dS* significantly above one, site 145. The sites shown in red are those that our geometrical model predicts to have *dN*/*dS* > 1. (Half of those sites have observed *dN*/*dS* > 1.) Note that our model predicts only sites on the basal side of sialic acid to be under positive selection, since our reference point for proximity is site 224. Site 145, the only purple site under positive selection, is also the only purple site on the basal side of sialic acid.

In summary, we have found that two simple geometric measures of a site’s location in the 3D protein structure, solvent exposure and proximity to the receptor-binding region, jointly outperformed, by a wide margin, any previously considered predictor of evolutionary variation in hemagglutinin, including immune epitope groups. In fact, the vast majority of the variation in evolutionary rate that was explained by the historical epitope sites was likely due to these sites simply being located near the receptor-binding region on the surface of the protein. However, historical epitope sites, in combination with solvent exposure and proximity, had some residual explanatory power beyond even a three-predictor model that combined the two geometric measures with experimental immune-epitope data. We suspect that this residual explanatory power reflects the sequence-based origin of the historical epitope sites. To our knowledge, the historical epitope sites were at least partially identified by observed sequence variation, so that, to some extent, these sites are simply the sites that have been observed to evolve rapidly in hemagglutinin.

## Discussion

We have conducted a thorough analysis of the determinants of site-specific hemagglutinin evolution. Most importantly, we have found that host immune bias (as currently measured by experimental and historical epitopes) accounts for a very small but significant portion of the evolutionary pressure on influenza hemagglutinin. In addition, we have found that epitope status cannot predict hemagglutinin sites under positive selection. By contrast, a simple geometric measure, receptor-binding proximity, is both a combined strong predictor of evolutionary rate and is the only quantity that can predict sites with *dN*/*dS* > 1. In addition, we have showed that a simple linear model containing three predictors, solvent accessibility, proximity to the receptor-binding region, and experimental epitopes, explains nearly 35% of the evolutionary rate variation in hemagglutinin H3. Therefore, our analysis suggests that one of two possible explanations must be true. First, it is possible that hemagglutinin antigenicity is not a strong direct driver of influenza adaptive evolution; rather, it is possible that influenza escapes the human host immune system by indirect means [23]. Second, alternatively, the current experimental data and historical epitopes may simply be insufficient and/or incorrect. Such a situation would explain why neither epitope definition can explain much evolutionary rate variation beyond the geometric constraints, and why neither epitope definition can predict sites under positive selection.

### History of epitopes in hemagglutinin H3

Efforts to define immune epitope sites in H3 hemagglutinin go back to the early 1980’s [10]. Initially, epitope sites were identified primarily by speculating about the chemical neutrality of amino acid substitutions between 1968 (the year H3N2 emerged) and 1977, though some limited experimental data on neutralizing antibodies was also considered [10, 11]. In 1981, the initial four epitope groups were defined by non-neutrality (amino-acid substitutions that the authors believed changed the chemical nature of the side chain) and relative location, and given the names A through D [10]. Since that original study in 1981, the names and general locations of H3 epitopes have remained largely unchanged [4,16]. The sites were slightly revised in 1987 by the same authors and an additional epitope named E was defined [11]. From that point forward until 1999 there were essentially no revisions to the codified epitope sites. In addition, while epitopes have since been redefined by adding or removing sites, no other epitope groups have been added [3,16,18]; epitopes are still named A–E. In 1999, the epitopes were redefined by more than doubling the total number of sites and expanding all of the epitope groups [4]. At that time, the redefinition consisted almost entirely of adding sites; very few sites were eliminated from the epitope groups. Although this set of sites and their groupings remain by far the most cited epitope sites, it is not particularly clear what data justified this definition. Moreover, when the immune epitope database (IEDB) summarized the publicly available data for influenza in 2007, it only included one experimental B cell epitope in humans (Table 2 in [36]). Although there were a substantial number of putative T cell epitopes in the database, *a priori* there is no reason to expect a T cell epitope to show preference to hemagglutinin as opposed to any other influenza protein; yet it is known that several other influenza proteins show almost no sites under positive selection. Moreover, it is known that the B cell response plays the biggest role is maintaining immunological memory to influenza, and thus it is the most important arm of the adaptive immune system for influenza to avoid.

The historical H3 epitope sites have played a crucial role in molecular evolution research. Since 1987, an enormous number of methods have been developed to analyze the molecular evolution of proteins, and specifically, to identify positive selection. The vast majority of these methods have either used hemagglutinin for testing, have used the epitopes for validation, or have at some point been applied to hemagglutinin. Most importantly, in all this work, the epitope definitions have been considered fixed. Most investigators simply conclude that their methods work as expected because they recover some portion of the epitope sites. Yet virtually all of these studies identify many sites that appear to be positively selected but are not part of the epitopes. Likewise, there is no single study that has ever found all of the epitope sites to be important. Even if the identified sites from all available studies were aggregated, we would likely not find every site among the historical epitopes in that aggregated set of sites.

### Implications of historical epitope groups for current research

Given all of this research activity, it seems that the meaning of an immune epitope has been muddled. Strictly speaking, an immune epitope is a site to which the immune system reacts. There is no *a priori* reason why an immune epitope needs to be under positive selection, needs to be a site that has some number or chemical type of amino acid substitutions, or needs to be predictive of influenza whole-genome or hemagglutinin-specific sequence cluster transitions. Yet, from the beginning of the effort to define hemagglutinin immune epitopes, such features have been used to identify epitope sites, resulting in a set of sites that may not accurately reflect the sites against which the human immune system produces antibodies.

Ironically, this methodological confusion has actually been largely beneficial to the field of hemagglutinin evolution. As our data indicate, if the field had been strict in its pursuit of immune epitopes sites, it would have been much harder to produce predictive models with those sites, in particular given that experimental data on non-linear epitopes have been sparse until very recently. By contrast, the historical epitope sites have been used quite successfully in several predictive models of the episodic nature of influenza sequence evolution. In fact, in our analysis, historical epitopes displayed the highest amount of variance explained among all individual predictors (Fig. 2). We argue here that the success of historical epitope sites likely stems from the fact that they were produced by disparate analyses each of which accounted for a different portion of the evolutionary pressures on hemagglutinin. Of course, it is important to realize that some of this success is likely the result of circular reasoning, since the sites themselves were identified at least partially from sequence analysis that included the clustered, episodic nature of influenza hemagglutinin sequence evolution.

Despite the success of historical epitope groups, they only predict about 18% of the evolutionary rate-variation of hemagglutinin for the entire phylogenetic tree. Since many of these sites likely are not true immune epitopes (and therefore not host dependent), one might ask which features of the historical epitope sites make them good predictors. We suspect that they perform well primarily because they are a collection of solvent-exposed sites near the sialic acid-binding region (see Fig. S2). We had shown previously that sites within 8 Å of the sialic acid-binding site are enriched in sites under positive selection, compared to the rest of the protein [20]. A similar result was found in the original paper by Bush et al. [4]. However, the related metric of distance from the sialic acid-binding site has not previously been considered as a predictor of evolution in hemagglutinin. Furthermore, before 1999, most researchers thought the opposite should be true; that receptor-binding sites should have depressed evolutionary rates [4]. Even today the field seems split on the matter [21]. As we have shown here, the inverse of the distance from sialic acid is a relatively strong quantitative predictor of hemagglutinin evolution; by itself this distance metric can account for 16% of evolutionary rate-variation. Moreover, by combining this one metric with another to control for solvent exposure, we can account for more than a third of the evolutionary rate variation in hemagglutinin. For reference, this number is larger than the variation one could predict by collecting and analyzing all of the hemagglutinin sequences that infect birds (another group of animals with large numbers of natural influenza infections), and using those rates to predict human influenza hemagglutinin evolutionary rates [20].

In terms of re-grouping experimental immune data, it is important to note that the IEDB has major limitations; not all existing (not to mention all possible) immunological data have been added. Further, the extent to which certain epitopes (e.g., stalk epitopes) have been mapped may be more reflective of a bias in research interests among influenza researchers than a bias in the human immune system. Also, until recently, the ability to generate unbiased high-affinity antibodies to influenza has been limited [37, 38]. Therefore, in our re-derivation of epitope groupings, we are certainly missing sites or may be incorrectly grouping the ones that we have. Our analysis of epitope sites will likely have to be redone as more data become available. However, we expect that as more non-linear data become available, they will broadly follow the trend observed in the linear epitope data, that is, the more antibodies are mapped, the more sites in the hemagglutinin protein appear in at least one mapping, until virtually every site in the entire hemagglutinin protein is represented. Under this scenario, the ability to predict evolution from immunological data would become worse, not better, as more data are accumulated.

One additional caveat comes from any potential effect of glycosylation on influenza immune escape. It is known that glycosylations on hemagglutinin can have a major effect on antibody binding [13]. In addition, the number of glycosylations in H3 hemagglutinin has increased since initial introduction of pandemic H3N2 in 1968 [13]. However, *a priori* there is no reason to believe that glycosylation will either increase or decrease *dN*/*dS* at individual sites or groups of sites; it could affect *dN*/*dS* in either direction, in particular if direct antibody escape is not the primary driver of hemagglutinin evolution. Moreover, there is no clear way to incorporate glycosylation into our regression model. In the future, investigating changing glycosylation patterns throughout the evolution of H3 hemagglutinin may yield important insights into influenza adaptation and immune escape.

### Geometric constraints likely dominate adaptive evolution in hemagglutinin

Why do geometric constraints (solvent exposure and proximity to receptor-binding site) do a good job predicting hemagglutinin evolutionary rates? Hemagglutinin falls into a class of proteins known collectively as viral spike glycoproteins (GP). In general, the function of these proteins is to bind a host receptor to initiate and carry out uptake or fusion with the host cell. Therefore, a priori one might expect that the receptor-binding region would be the most conserved part of the protein, since binding is required for viral entry. Yet, in hemagglutinin sites near the binding region are the most variable in the entire protein. There are at least two possible models that might explain this observation. First, conventional wisdom says that in terms of host immune evasion, antibodies that bind near the receptor-binding region may be the most inhibitory, and hence mutations in this region the most effective in allowing immune escape. Viral spike GPs have a surface that is both critical for viral survival and is sufficiently long lived that a host immune response is easily generated against it. There are likely many other viral protein surfaces that are comparatively less important or sufficiently short lived during a conformational change that antibody neutralization is impractical. Thus, the virions that survive to the next generation are those with substantial variation at the surface or surfaces with high fitness consequences and a long half-life in vivo. Evolutionary variation at surfaces with low or no fitness consequences, or at short-lived surfaces, should behave mostly like neutral variation and hence appear as random noise, not producing a consistent signal of positive selection. Second, according to the avidity modulation model of Hensley et al. [23], it is possible that antibody inhibition is not overcome by escaping the antibody directly. Rather, a single or a few relatively rare mutations may increase the avidity of hemagglutinin for its receptor so as to out-compete partial antibody inhibition. Subsequently, once the partial inhibition is overcome in a competent host, passage to an incompetent host allows genetic drift to bring the avidity back down to baseline. Considering the fact that neither historical nor experimental immune epitopes vastly out-performed our simple distance metric, we think that our results support the avidity modulation model [23], which does not predict a bias based on antibody binding sites. However, it remains a possibility that the historical epitopes and current experimental data are simply wrong about which sites and groups of sites the human immune system attacks. Either way, our work highlights the need for a paradigm shift in the field.

We also need to consider that actual epitope sites, i.e., sites toward which the immune system has a bias, may not be that important for the evolution of viruses. An epitope is simply a part of a viral protein to which the immune system reacts. Therefore, it represents a host-centered biological bias. The virus may experience stronger selection at regions with high fitness consequences but that generate a relatively moderate host response compared to other sites with low fitness consequences that generate a relatively strong host response. Moreover, there is little reason to believe that influenza *must* escape an antibody by directly reducing the binding of that antibody. There are many other possible scenarios for immune evasion. Thus, we expect that the geometric constraints we have identified here will be more useful in future modeling work than the experimental epitope groups we have defined. Moreover, we expect that similar geometrical constraints will exist in other viral spike glycoproteins, and in particular in other hemagglutinin variants.

By contrast to the clear geometric constraints we observed for the pre-fusion structure, we found no comparable result for the post-fusion structure. There are perhaps several good reasons to expect this result. First, the transition state is likely very short-lived, such that the human immune system is not able to generate antibodies against it. Second, due to the short-lived functional nature of the transition state, there is likely relatively little selection for folding stability. Therefore, for the post-fusion structure we do not expect to observe the RSA–rate correlation that exists in the pre-fusion structure and in most other proteins. Third, models describing the transition from the pre-fusion to the post-fusion state show that the HA1 chain dissociates from the HA2 chain [39]. Subsequently, the HA2 chain carries out virtually all of the fusogenic functions. Thus, the HA1 chain is likely the functional unit in the first step of entry and the HA2 chain is likely the functional unit in the second. However, there is almost no rapid evolution happening in the HA2 chain, i.e., the HA2 chain does not seem to experience any positive diversifying selection.

Remarkably, the sites we found that experienced the most positive selection showed minimal overlap with the sites found to be minimally sufficient for explaining the major antigenic transitions in H3N2, as determined by HI assays with ferret antisera [21]. While both groups of sites lie near the sialic-acid binding region, the vast majority of positively selected sites are located basally to sialic acid whereas sites identified by HI assays lie predominantly on the apical side (Fig. 4). This finding suggests that HI assays and positive selection analyses reflect distinct biological mechanisms. For example, HI assays might not accurately reflect selection pressures *in vivo*. Such a result would suggest that influenza is not under pressure to directly escape antibody binding. Alternatively, HI assays may correctly identify mutations that lead to antigenic cluster transitions whereas positive selection analyses may identify sites that mediate avidity [23] or antigenic drift within a cluster. Yet another alternative is that the standard manner for obtaining ferret antisera simply may not represent a good proxy for the cyclical nature of human influenza infections [40]. Indeed, recent evidence suggests that, at least for the pandemic H1N1 strain, cyclical infections can shift the antibody response toward the receptor-binding region [41]. In future work, disentangling the different mechanisms reflected by HI assays and by positive-selection analyses will likely be crucial for improved prediction of HA evolution and of optimal vaccine strains.

## Materials and Methods

### Obtaining influenza data and preparing sequences

All of the data we analyzed were taken from the Influenza Research Database (IRD) [42]. The IRD provides experimental immune epitope data curated from the data available in the Immune Epitope Database (IEDB) [43].

We used sequences that had been collected since the 1991–1992 influenza season. Any season before the 1991–1992 season had an insufficient number of sequences to contribute much to the selection analysis. The sequences were filtered to remove redundant sequences and laboratory strains. The sequences were then aligned with MAFFT [44]. Since it is known that there have been no insertions or deletions since the introduction of the H3N2 strain, we imposed a strict opening penalty and removed any sequences that had intragenic gaps. In addition, we manually curated the entire set to remove any sequence that obviously did not align to the vast majority of the set; in total the final step only removed about 10 sequences from the final set of 3854 sequences. For the subsequent evolutionary rate calculations, we built a tree with FastTree 2.0 [45] .

### Computing evolutionary rates and relative solvent accessibilities

To compute evolutionary rates, we used a fixed effects likelihood (FEL) approach with the MG94 substitution model [24,46,47]. We used the FEL provided with the HyPhy package [24]. For the full setup see the linked GitHub repository (https://github.com/wilkelab/influenza_HA_evolution). As is the case for all FEL models, an independent evolutionary rate is fit to each site using only the data from that column of the alignment. Because our data set consisted of nearly 4000 sequences, almost every site in our alignment had a statistically significant posterior probability of being either positively or negatively selected after adjusting via the false discovery rate (FDR) method. As shown in Figure 3, all evolutionary rates fall into a range between *dN*/*dS* = 0 and *dN*/*dS* = 4.

We computed RSA values as described previously [28]. Briefly, we used DSSP [48] to compute the solvent accessibility of each amino acid in the hemagglutinin protein. Then, we used the maximum solvent accessibilities [49] for each amino acid to normalized the solvent accessibilities to relative values between 0 and 1. We found that RSA calculated in the trimeric state produced better predictions than RSA calculated in the monomeric state. Thus, we used multimeric RSA in all models in this study. Both multimeric and monomeric RSA are included in the supplementary data.

### Evolutionary rate-distance correlations

To create the structural heat map of correlations shown in Fig. 1B, we first needed to calculate the correlations between evolutionary rates and pairwise distances, calculated in turn for each location in the protein structure as the reference point for the distance calculations. Conceptually, we can think of this analysis as overlaying a grid on the entire protein structure, where we first calculate the distance to various grid points from every *C_α_* in the entire protein, and then compute the correlation between the set of distances to the sites on the grid and the evolutionary rate at those sites. In practice, we calculated the distance from each *C_α_* to every other *C_α_*. We then colored each residue by the correlation obtained between evolutionary rates and all distances to its *C_α_*.

### Statistical analysis and data availability

All statistical analyses were performed using R [50]. We built the linear models with both the `lm()` and `glm()` functions. For cross validation, we used the `cv.glm()` function within the boot package. Residual standard error values were computed by taking the square root of the delta value from `cv.glm()`. With the exception of graph visualizations, all figures in this manuscript were created using ggplot2 [51].

A complete data set including evolutionary rates, epitope assignments, RSA, and proximity to the receptor-binding site is available as Table S1. Raw data and analysis scripts are available at https://github.com/wilkelab/influenza_HA_evolution. In the repository, we have included all human H3 sequences from the 1991–1992 season to present combined into a single alignment. We have cleaned the combined data to only include sequences with canonical bases, non-repetitive sequences, and we have hand filtered the data to ensure all included sequences align appropriately to the 566 known amino acid sites. In addition, we have built a tree and visually verified that there were no outlying sequences on the tree for the combined set.

### Technical considerations for analysis

The site-wise numbering for the H3 hemagglutinin protein reflects the numbering of the mature protein; this numbering scheme requires the removal of the first 16 amino acids in the full-length gene. Thus, for protein numbering purposes, site number 1 is actually the 17th codon in full-length gene numbering. The complete length of the H3 hemagglutinin gene is 566 sites while the total length of the protein is 550 sites. It is important to point out that the mature H3 protein has two chains (HA1 and HA2) that are produced by cutting the presursor (HA0) protein between sites 329 and 330 in protein numbering. In addition, as a result of cloning and experimental diffraction limitations, most (or likely all) hemagglutinin structures do not include some portion of the first or last few amino acids of either chain of the mature protein, and crystallographers always remove the C-terminal transmembrane span from HA2. For example, the structure we used (PDBID: 4FNK) in this study does not include the first 8 amino acids of HA1, the last 3 amino acids of HA1, or the last 48 amino acids of HA2. As a result, HA1 includes sites 9–326 and HA2 includes sites 330–502. The complete data table in the project repository lists the gene sequence from one of the three original H3N2 (Hong Kong flu) hemagglutinin (A/Aichi/2/1968), the gene numbering, the protein numbering, the numbering of one H3N2 crystal structure, historical immune epitope sites from 1981, 1987 and 1999, and every calculated parameter used (and many others than were not used) in this study. In general, the most common epitope definitions in use today are those employed by Bush et. al 1999 [4]. Throughout this work, we refer to the Bush et. al 1999 epitopes as the “historical epitope sites”.

## Acknowledgments

We would like to thank Jesse Bloom and Trevor Bedford for helpful comments on this manuscript and Robin Bush for providing us with a complete list of the historical epitope groupings.

## Supporting Information Legends

**Data Table S1: Complete data set including evolutionary rates, solvent accessibilities, proximities to the receptor-binding region, and epitope status for all sites.**

**Text S1: Analysis of available experimental human epitope data.**

